# Developing a xenograft model of human vasculature in the mouse ear pinna

**DOI:** 10.1101/789537

**Authors:** Gavin R. Meehan, Hannah E. Scales, Rowland Osii, Mariana De Niz, Jennifer C. Lawton, Matthias Marti, Paul Garside, Alister Craig, James M. Brewer

## Abstract

Humanised xenograft models allow for the analysis of human tissue within a physiological environment *in vivo*. However, current models often rely on the angiogenesis and ingrowth of recipient vasculature to perfuse tissues, preventing analysis of biological processes and diseases involving human blood vessels. This limits the effectiveness of xenografts in replicating human physiology and may lead to issues with translating findings into human research.

We have designed a xenograft model of human vasculature to address this issue. Human subcutaneous fat was cultured *in vitro* to promote blood vessel outgrowth prior to implantation into immunocompromised mice. We demonstrate that implants survived, retained human vasculature and anastomosed with the circulatory system of the recipient mouse. Significantly, by performing transplants into the ear pinna, this system enabled intravital observation of xenografts by multiphoton microscopy, allowing us to visualise the steps leading to vascular cytoadherence of erythrocytes infected with the human parasite *Plasmodium falciparum*.

This model represents a useful tool for imaging the interactions that occur within human tissues *in vivo* and permits visualization of blood flow and cellular recruitment in a system which is amenable to intervention for various studies in basic biology together with drug evaluation and mechanism of action studies.

## Introduction

Xenograft models, in which human cells, tissues or organs are implanted into immunodeficient animal hosts have proven to be valuable research tools in recent years^1,2^. By allowing human tissue to be studied in an *in vivo* environment, these models provide a physiologically relevant method for assessing human diseases that neither conventional animal models nor human *in vitro* studies are able to achieve alone.

A wide array of tissues have been examined in these models including hematopoietic stem cells, liver and thymus fragments^3^, adipose tissue^4^, testicular tissue^5^, tumours^6^ and even human brain organoids^7^. However, the majority of xenografts tend to consist of cells or small organoids as a major challenge of using tissue fragments is the re-establishment of blood flow to the grafts. When a tissue is first implanted into an animal, cells within a distance of 150-200µm of a blood vessel survive through molecular diffusion but those cells deeper in the graft experience hypoxia and glucose deprivation^8^. This triggers the release of soluble mediators including vascular endothelial growth factor (VEGF) which induce both host and graft endothelial cells (ECs) to initiate angiogenesis^9^. This allows the ECs to self-replicate and form hollow capillary sprouts that continue to grow until they meet and connect with another capillary, resulting in the restoration of normal blood flow.

During this process of neovascularisation, the vessels from the host tend to grow into the graft i n a process known as internal inosculation. This results in a substantial proportion of the graft vasculature originating from the host, and a loss of the vasculature of graft origin^10,11^, limiting the ability to effectively study the function of human vasculature within human xenografts^12^.

Research in a variety of areas would benefit from a xenograft system that would allow non-invasive *in vivo* visualisation of cellular interactions occurring within and around human vasculature. These include research examining vascular remodelling and repair, models of acute inflammation as well as various infection studies examining the interactions of bacteria or protozoan parasites with human tissue^13^. A model allowing for the visualisation of the vascular cytoadherence of *Plasmodium falciparum* would be particularly useful, as this process is thought to be a major contributor to a deadly form of disease^14^, known as cerebral malaria, but cannot be replicated in murine malaria models.

Accordingly, we designed a new model that incorporated human tissue in a site that was minimally invasive and allowed for longitudinal imaging; the ear pinna. We demonstrated that by pre-culturing the tissue prior to implantation we could promote the retention of human vasculature and the successful re-perfusion of the graft. Using the human specific parasite *Plasmodium falciparum*, we validated the function of the model and visualised binding of the parasite to the endothelium.

## Methods

### Mice

NOD *SCID* Gamma mice (NSG) ^15^ (originally purchased from Jackson Laboratories, Bar Harbor, ME, USA) were produced in house (Central Research Facility, University of Glasgow, UK). These mice were bred in a sterile film isolator and maintained in individually ventilated cages (IVC). Animals were maintained on a 12-hour light/dark cycle and provided with food and water *ad libitum*. All procedures were performed under a UK Home Office licence in accordance with the Animals (Scientific Procedures) Act 1986.

### Ethical Approval

The use of human tissue within this study was approved by the NHS Greater Glasgow and Clyde Biorepository (application number 97) on behalf of the NHS Research Ethics Committee. Informed consent was obtained from all subjects. All tissue was processed in accordance with the Human Tissue (Scotland) Act 2006.

### Human Tissue

Human tissue was acquired from patients undergoing routine surgery at the Queen Elizabeth University Hospital (QEUH), Glasgow, UK. Tissue was collected by the Greater Glasgow and Clyde Biorepository on our behalf. The tissue consisted of subcutaneous fat from the breast or abdomen of female patients between the ages of 20 and 60 years old. Tissue was only taken from patients not receiving ongoing cancer treatment.

### Tissue Culture

Human subcutaneous fat was stored in Aqix (Life Science Production, UK) prior to culture. Under aseptic conditions, small pieces of tissue (3-5mm^2^) were dissected and 4 pieces placed in each well of a 12 well plate. Matrigel (400μl; Corning, NY, USA) was then added to each well and the plate was incubated at 37°C for 30 minutes to polymerise the Matrigel. The wells were then supplemented with 2ml of endothelial growth medium (EGM) (Promocell, Germany). The plates were incubated at 37°C in 5% CO2. Media was replaced every three days.

### Ear Implant Surgery

After 7 days of culture, tissue with more than 0.25mm of external inosculation around the entire outer circumference was implanted into mice (Figure 1E). All surgical procedures took place in a Category 2 biological safety cabinet under aseptic conditions. The mice were anaesthetised with oxygen (1.5L/minute) containing 2.5% isoflurane and prophylactically treated with buprenorphine (0.1mg/kg). The ear was cleaned with 70% alcohol. It was then affixed to double sided adhesive tape on the bore of a 5ml syringe. The ear pinna was injected with 50-100μl of sterile saline to create a pocket. Forceps were then inserted into the ear to create an incision, which was enlarged using microdissection scissors. Curved forceps were used to enlarge the pocket and separate the skin of the ear pinna. The Matrigel embedded tissue was then inserted into the pocket and the incision was sealed using Vetbond glue (3M, MN, USA). Mice were recovered with oxygen before being moved to an IVC placed on a heat mat for 1 hour. They were returned to normal housing and monitored daily until the implants had fully healed. All experiments were carried out using these mice 21-35 days post-surgery unless otherwise stated.

**Figure 1.**
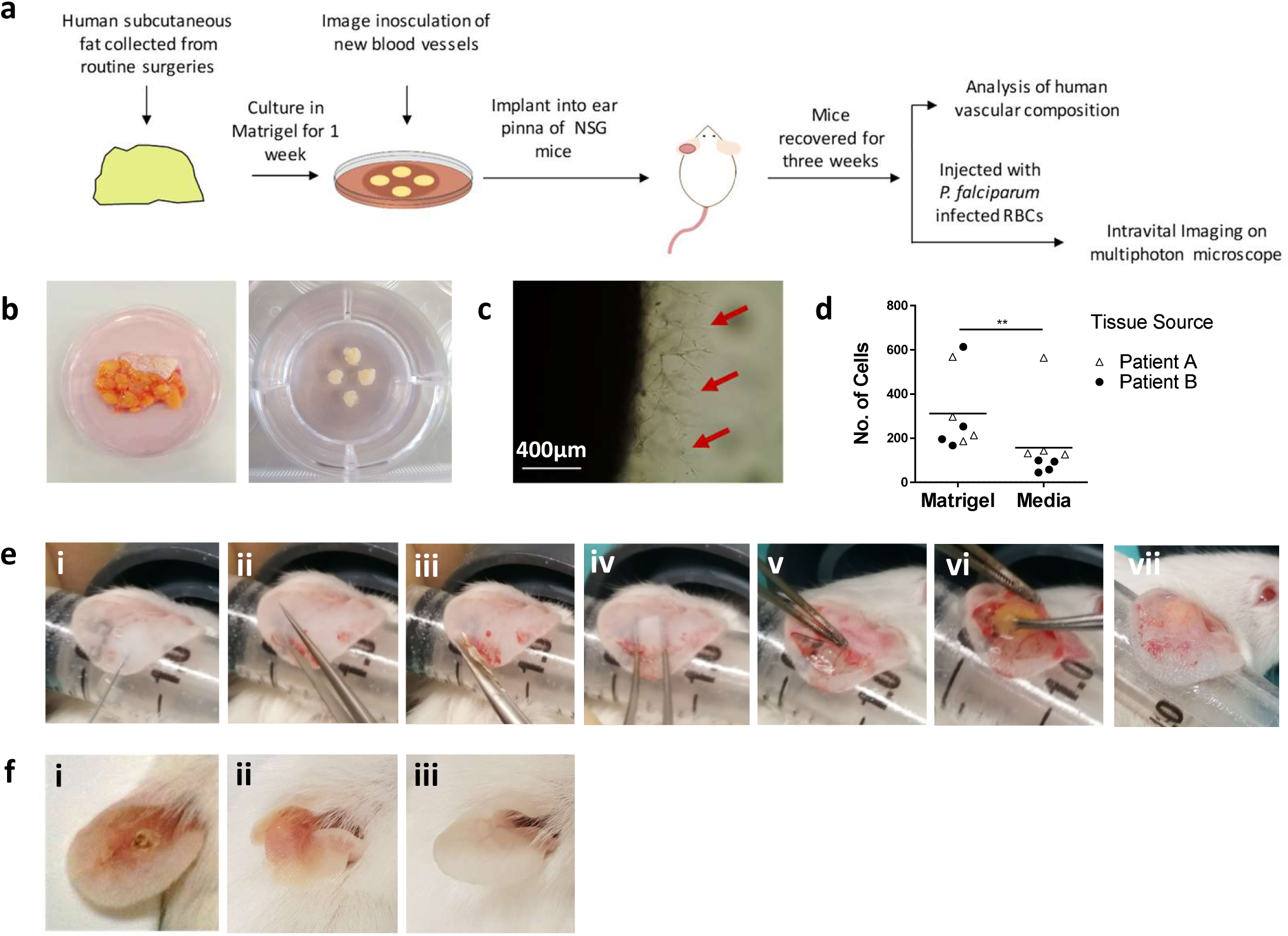
Developing a method for implanting human tissue into the ear pinna of immunocompromised mice. (a) Schematic of the model. (b) Human subcutaneous fat was collected from routine surgeries and dissected into 3-5mm^2^ sized pieces before being embedded in Matrigel (c) which promoted the outgrowth of blood vessels (red arrows). (d) Comparisons of Matrigel embedded versus non Matrigel embedded tissues indicated an increase in the number of endothelial cells in embedded tissues. (e) The surgery was performed by (i) affixing the ear pinna to a syringe and injecting 50µl of saline to create a pocket. (ii) A hole was created using forceps. (iii) Scissors were inserted in this hole and used to create a flap. (iv) Curved forceps were used to split the skin apart to create a larger pocket (v) The superior skin was retracted and (vi) the human tissue was inserted. (vii) The incision was sealed with glue and the mouse recovered. (f) (i) Ears implanted with human tissue using a previous method^23^ did not heal well. (ii) Ears implanted with tissue using the new method healed well three weeks post-surgery and resembled the (iii) control ear. (d) n=8, pooled from two independent experiments. Statistical analysis was performed using a Mann Whitney test, **p<0.01.

### Flow Cytometry

Comparisons of fresh, cultured and implanted tissues were performed using flow cytometry. The tissues were all processed using the same method with some minor changes. Prior to digestion, each well containing cultured tissue was incubated with 1mg/ml dispase II (Thermofisher, Waltham, MA, USA) in Ca^2+^/Mg^2+^ free PBS for 1 hour at 37°C to dissolve the Matrigel. The tissue in these wells were analysed together and the results divided by the number of embedded pieces of tissue in the well. Engrafted mice were culled 21 days post-surgery and their implant and control ears removed and placed in RPMI prior to digestion. All other processing steps were identical between the different tissue conditions.

Once collected, the tissues were minced and digested in RPMI containing 2% foetal bovine serum (FBS) (Thermofisher, Waltham, MA, USA), 1mg/ml collagenase II ((Sigma Aldrich, Dorset, UK) and 10U/ml DNase (Invitrogen, Carslbad, CA, USA) for 50 minutes at 37°C under constant agitation. The tissues were processed through a 100µm filter and washed twice in 2% RPMI and once in Ca^2+^/Mg^2+^ free PBS containing 2mM EDTA at 400*g* for 5 minutes. Blood was not digested. The cell solutions were then suspended in 400µl Ca^2+^/Mg^2+^ free PBS containing e520 viability dye (1:1000) (eBiosciences, Waltham, MA, USA) and incubated for 20 minutes at 4°C. This step was omitted in samples examining labelled infected red blood cells. The cells were washed in Ca^2+^/Mg^2+^ free PBS containing 2mM EDTA and resuspended in conditioned media from the anti-CD16/CD32 antibody producing hybridoma (2.4G2) (FcBlock). To examine human endothelial cells, antibody suspensions containing human Fc Block, V450 labelled anti-human CD36 (both from BD Biosciences, San Jose, USA), APC-e780 labelled anti-human CD45, APC labelled anti-mouse CD31, PE labelled anti-human CD31, AF700 labelled anti-human CD34 (all from Biolegend, San Diego, CA, USA), PE-Cy7 labelled anti-mouse CD45 and PerCP-e710 labelled anti-human ICAM-1 (both from eBiosciences, Waltham, MA, USA) were added to each sample and incubated for 20 minutes at 4°C. To examine labelled infected red blood cells *in vivo*, antibody suspensions were prepared using PE-Cy7 labelled anti-mouse CD45 as above, APC labelled anti-mouse Ter119 (from eBiosciences, Waltham, MA, USA) and, depending upon the experiment, PerCP-Cy5.5 labelled anti-human CD235a (from Biolegend, San Diego, CA, USA) and incubated as described above. The cells were subsequently washed and analysed using a BD LSRfortessa flow cytometer (BD Biosciences, San Jose, USA). Analysis of flow cytometry data was performed using FlowJo 10 (FlowJo LLC, Ashland, OR, USA).

### Histology

Mice were euthanized, and both the implant and controls ears were cut off and placed in 10% neutral buffered formalin for 16 hours. The tissue was then dehydrated through different grades of alcohol and xylene and embedded in wax. 8μm transverse sections were cut from each tissue and mounted on glass slides. They were then stained with haematoxylin and eosin and subsequently imaged on an EVOS Cell Imaging System (Thermofisher, Waltham, MA, USA).

### Immunofluorescence

Mice were euthanized, and both the implant and controls ears were cut off and placed in PBS. Using forceps, the skin on the ventral side of the ear was retracted to expose the human tissue implant. The ears were fixed in 4% paraformaldehyde for 20 minutes at 4°C and incubated overnight in 0.05% Triton X-100 in PBS containing anti-human CD31 (Biolegend, San Diego, CA, USA) antibody conjugated to either PE or AF647 in a 1:200 dilution. The tissue was washed in PBS and mounted on a glass slide with a glass coverslip for imaging.

### Parasites

*Plasmodium falciparum* was cultured as described previously^16,17^. Cultures were grown in RPMI-1640 supplemented with 24mM sodium bicarbonate (Sigma-Aldrich, St Louis, MO, USA) and 10% human serum (Interstate Blood Bank Inc, Memphis, TN, USA). They were gassed with 5% CO2/1% O2 and 94% N2 mixture and maintained at 37°C. The ICAM-1/CD36 binding parasite strain ItG was used throughout^18^.

Parasites were magnetically purified using CS columns (Miltenyi-Biotech, Bergisch Gladbach, Germany) in a SuperMACS II separator (Miltenyi-Biotech, Bergisch Gladbach, Germany) as per manufacturer’s instructions. Columns were blocked with 3% BSA for 15 minutes. One 50mmx9mm petri dish containing 25ml of parasites, maintained at 8-10% parasitaemia at a 5% haematocrit produced approximately 6×10^8^ infected RBCs at a purity of 80-90%. The parasites were ready to use when in mid/late trophozoites stages. They were washed and resuspended in 20ml of serum free RPMI and added to the column. Once the parasites had run through the column, it was washed with 25ml of serum free RPMI. The column was detached from the magnet and the parasites were removed by washing with 50ml of serum free RPMI.

### Labelling red blood cells

Red blood cells (RBCs) with and without parasites were fluorescently labelled with CMTPX (Thermofisher, Waltham, MA, USA). All labelling and incubation steps took place at 37°C under constant agitation. RBCs were incubated with CMTPX (5μM) for 20 minutes. 1ml of dye was used to stain 1×10^8^ RBCs. Following staining, the RBCs were washed twice at x400g for 5 minutes and incubated with 10% RPMI for 10 minutes. The RBCs were washed a further two times at x400g for 5 minutes in Ca^2+^/Mg^2+^ free PBS and resuspended in 100µl of Ca^2+^/Mg^2+^ free PBS.

### Static binding assay

On a marked plastic 60mm × 15mm petri dish, 2µl drops of human ICAM-1 (R&D Systems, MN, USA), mouse ICAM-1 (R&D Systems, MN, USA) or PBS were added in triplicate around the sides of the dish. The dish was placed in a humidity chamber and incubated at 37°C for 1 hour. The spots were then removed, and the dish was blocked in 1% BSA in PBS at 37°C for 1 hour. A suspension of *P. falciparum* trophozoites at 3% parasitaemia in 1% haematocrit was prepared. 1.25ml of parasite suspension was added to the dish, which was swirled occasionally, for 1 hour at 37°C. The parasites were gently removed, and the dish was gently washed in RPMI 5 times. The bound parasites were then fixed in 1% glutaraldehyde in PBS at room temperature for 20 minutes. Imaging of the dish was performed using an EVOS Cell Imaging System (Thermofisher, Waltham, MA, USA). One image was taken in the centre of each spot and the number of bound parasites was calculated.

### Multiphoton Imaging of Implants

Multiphoton imaging was performed with a Zeiss LSM7 MP system equipped with both a 10x/0.3 NA air and a 20x/1.0NA water immersion objective lens (Zeiss) and a tuneable titanium/sapphire solid state 2-photon excitation source (Chamelon Ultra II; Coherent Laser Group) and optical parametric oscillator (OPO; Coherent Laser Group). For intravital imaging, mice were given an intravenous injection of 20µg of PE labelled or AF647 labelled anti-human CD31 and 5µg TNFα in PBS. Following six hours of incubation, the mice were injected intravenously with 5×10^8^ uninfected unlabelled RBCs and anaesthetised with oxygen (1.5L/minute) containing 2.5% isoflurane. The implant or control ear was affixed to a metal stage using veterinary-grade glue (Vetbond; 3M). Silicone grease was used to create a ring around the ear, which was then filled with PBS (Supplementary Figure 4). When under the microscope, mice were given an injection of 5×10^8^ CMTPX labelled infected red blood cells in the tail vein. A laser output of 820 nm with or without an OPO signal at 1060 nm provided excitation of adipocytes, hCD31 positive cells and labelled red blood cells. Images were acquired with an X-Y pixel resolution of 512 × 512 in a variety of Z increments. For vessel analysis, 5×5 tile scans of 100µm stacks were acquired in 4µm Z stack increments. All images were processed using Volocity 6.1.1 (Perkin Elmer, Cambridge, UK)

### Image Analysis

Analysis of human blood vessels was performed using images of fresh and implanted human tissue from different patients. Between 2 and 3 animals implanted with the same human tissue were used per data point for the implanted images. All images were analysed using the Vessel Analysis plugin provided by Fiji. Z stacks were first flattened then converted to binary images. The vascular density plug in was then run to produce the vascular density and vascular length density measurements. Vessel diameters were measured manually by counting five randomly selected vessels in five fields of view in each image.

Analysis of infected red blood cells *in vivo* was performed using Imaris Version 7.6.5. Individual cells were defined as objects and tracked in 2D. Track plots demonstrate the movement of cells relative to their point of origin in the blood vessels.

### Statistical Analysis

All graphs and statistical analyses were produced using GraphPad Prism 7 (GraphPad Software Inc., San Diego, CA, USA). P values <0.05 were deemed to be significant.

## Results

### Developing a method for implanting human tissue into the ear pinnae

The basic layout of the xenograft model is shown in Figure 1A. We focussed specifically on subcutaneous adipose tissue (SAT) for development of the model as it is well vascularised and is an important site in both infection and immunity^19,20^. The tissue is also a common by-product of surgery and was therefore readily available for technical optimisation. Although not characterised as fully as SAT, we also had limited success implanting dermis, kidney cortex and skeletal muscle into the ear pinna as shown from histology studies of the tissues three weeks post-implantation. However, we were unable to form stable grafts consisting of brain, duodenum or colon as these tissues underwent major morphological changes upon implantation (Figure S1).

Initial work examined whether tissue could be implanted directly from patients into the ear pinna of mice. We found that these tissues healed poorly and lost a substantial proportion of graft vasculature (data not shown). We therefore decided to culture the human tissue prior to implantation to stimulate the outgrowth of blood vessels^21^. To achieve this, tissues were embedded in Matrigel supplemented with endothelial growth medium (Figure 1B). Consistent with previous studies^22^, this improved the retention of vessels of graft origin (Figure 1C). Flow cytometric analysis of engrafted implants indicated that a significantly higher number of human ECs (Mann-Whitney, P<0.01) were found in Matrigel embedded tissues compared to those cultured without Matrigel (Figure 1D). This confirmed that the pre-treatment improved the retention of the human vasculature in grafts, as shown in previous animal engraftment studies^22^.

Following 7 days of culture, tissues with externally inosculated vessels at least 0.15mm in length around their entire outer circumference were implanted into NSG mice. Our previously described implantation method^23^ resulted in poor wound healing with human tissue (Figure 1Fi). We therefore optimised the surgical approach by creating a large pocket in the mouse ear pinna (Figure E). To achieve this, sterile saline was first injected into the ear pinna to create a subcutaneous pouch (i). Forceps were then placed into the needle entry site to enlarge the opening (ii) and an incision was made using scissors (iii). Curved forceps were then used to expand the pouch and separate the skin (iv). The skin on the ventral side was retracted (v) and Matrigel embedded SAT was placed within the pouch (vi). Finally, the incision was sealed using veterinary grade glue (vii). Following three weeks of recovery, the ear pinna had fully healed (Figure 1Fii) and resembled the control ear (Figure 1Fii) confirming that tissue could be successfully engrafted using this method.

### Human tissue survives and retains human vasculature following implantation into the ear pinna of immunocompromised mice

Having developed a method for implanting human tissue into the ear pinna, we next had to determine whether the grafts survived and retained human vasculature. Histological analysis of implant (Figure 2A) and control (Figure 2B) ears demonstrated that the adipocytes and stroma of the transplanted tissue retained their normal structural characteristics and did not form a necrotic core. The grafts survived for up to five weeks post-implantation but may survive longer as has been shown in previous human engraftment studies^4,5^. Flow cytometry analysis was performed to identify transplant ECs (mouse CD45-(mCD45)/human CD45-(hCD45), mouse CD31-(mCD31)/human CD31+ (hCD31)) in the fresh, cultured and implanted human SAT (Figure S2). Examination of the total number of human ECs indicated that the implants contained a significantly higher number of ECs than the fresh tissues (Figure 2C) (Kruskal-Wallis test, P<0.05). A breakdown of the EC subpopulations indicated few changes in the proportions of ECs expressing the adhesion molecules CD34 (Figure S3A) and ICAM1 (Figure S3B) suggesting that the cells had not become activated as a result of implantation. However, the percentage of ECs expressing the scavenger marker CD36 (Figure S3C) was significantly lower in implanted tissue suggesting a transplant induced reduction in the subcutaneous microvasculature characteristics of the tissue^24^ (Two Way ANOVA, P<0.0001).

**Figure 2.**
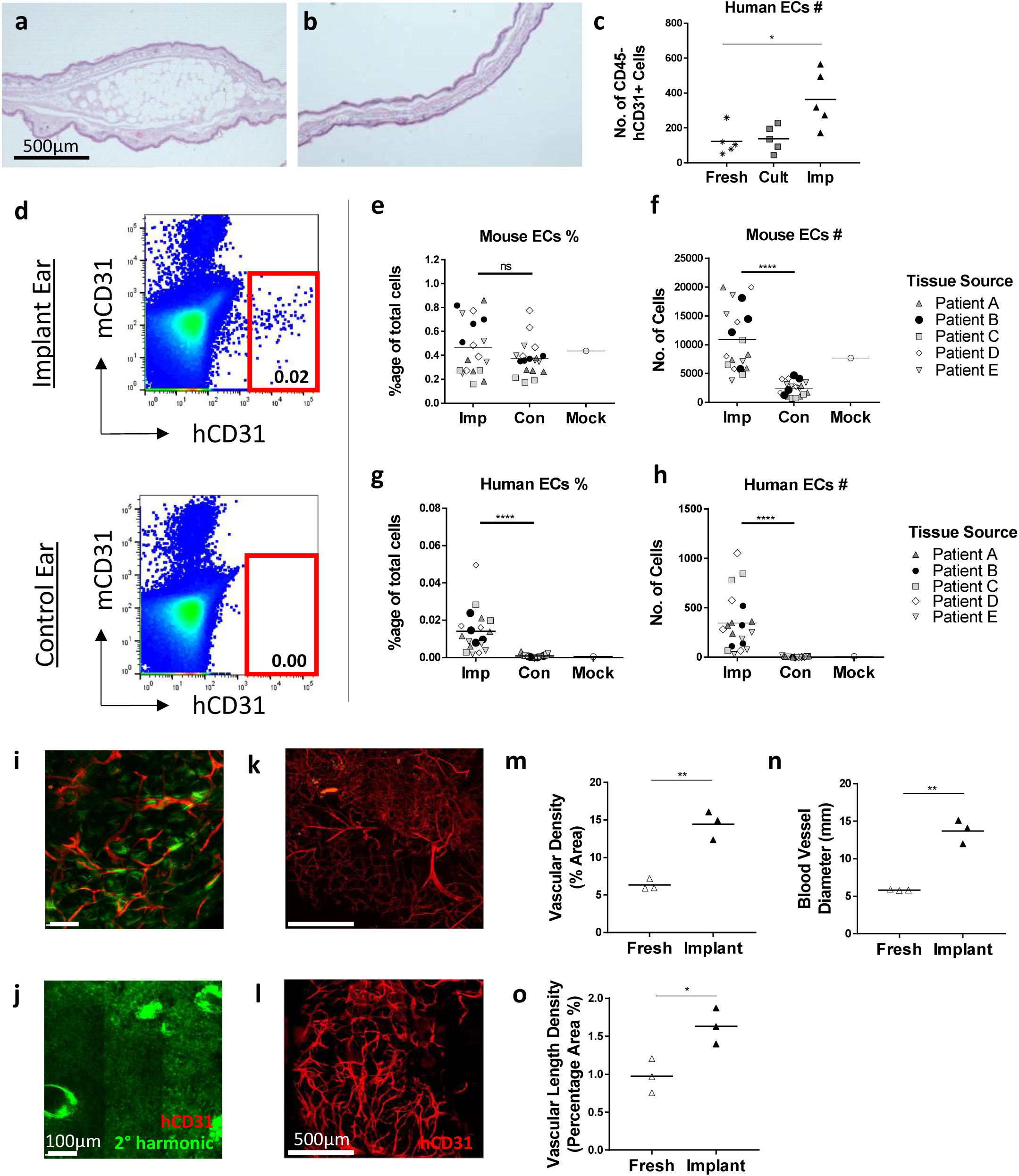
Human vasculature is retained in engrafted tissue. Analysis was performed on fresh, cultured and implanted human tissue to determine the effect of engraftment. Histological staining of the (a) implant and (b) control ear demonstrated that the human tissue survived implantation. (c) Flow cytometry was performed to determine the endothelial cell (EC) content of fresh, cultured (Cult) and implanted (Imp) tissue. n=5 from five independent experiments. Each data point from implanted samples was pooled from 3-4 mice implanted with human tissue from the same source. Statistical analysis was performed using a Kruskal-Wallis test, *p<0.05. (d-h) Flow cytometry was performed on implant (Imp), mock surgery (Mock) and control (Con) ears to examine the mouse and human ECs populations. 3-4 mice per group, five independent experiments. Statistical analysis was performed using a Wilcoxon test, ****p<0.00 01. (i) An intravenous injection of an anti-human CD31 antibody confirmed that the engrafted tissue contained human ECs and was perfused. (j) The control ear confirmed that the antibody w as human specific. Tile scans were taken of (k) fresh and (l) implanted human tissue stained with an anti-human CD31 antibody. Images were centred on the core of the tissues but included most vessels on the periphery. The changes in the blood vessels were quantitatively measured using a Fiji plugin called Vessel Analysis which determined the (m) vascular density, and (n) vascular length density. Blood vessel diameters (o) were measured manually by counting five randomly selected blood vessels in five fields of view in each image. n=3 from three independent experiments. Each data point from implanted samples was pooled from 2-3 mice implanted with human tissue from the same source. Statistical analysis was performed using an unpaired T test, *p<0.05, **p<0.01.

Flow cytometry of control, mock surgery (surgery with no tissue implants) and implant ears indicated the presence of human ECs in the implant ears only (Figure 2D). The percentage of mouse EC’s was not significantly different between treatments (Figure 2E) but the number of mouse EC’s was significantly higher in the implant ear compared to the control ear (Figure 2F) (Wilcoxon Test, P<0.0001). The number of cells were similar to those observed in the mock surgery ear suggesting that the increase was related to healing in response to surgery. By contrast, the percentage (Figure 2G) and number (Figure 2H) of human ECs were found to be significantly increased in the implant ears compared to the control ears (Wilcoxon Test, P<0.0001) and were completely absent in the mock surgery control. This confirmed that the human ECs had been retained and had expanded in the implant ear following surgery.

To definitively confirm that the tissues had successfully anastomosed with the recipient mouse vasculature we intravenously injected engrafted mice with a human specific αCD31 antibody. After one hour, we culled the mouse and imaged the tissue using multiphoton microscopy. Analysis of these images clearly demonstrated that the implanted tissues contained human blood vessels (Figure 2I) whereas no human vessels were detected in the mouse control ears (Figure 2J).

Further analysis of the vasculature of hCD31 labelled fresh (Figure 2K) and implanted (Figure 2L) tissues indicated significantly higher vascular densities (Figure 2M) and vascular length densities (Figure 2O) in the implanted tissue (Unpaired T test, P<0.05) particularly around the periphery of the graft (Figure S4), as per previous engraftment research^25^. Similarly, the blood vessel diameters (Figure 2N) were also found to be significantly larger in the implanted tissue suggesting a reduction in the microvasculature as shown by flow cytometry (Figure S3C). The human vasculature was present for up to five weeks in our study. Although we did not examine later time points we would expect it to survive longer as shown with previous human EC engraftment studies^26^.

### Tracking labelled red blood cells *in vivo*

To image the intravascular behaviours of *P. falciparum* to human endothelium, we selected an ICAM-1 binding parasite known as ItG^18^. From our flow cytometry data (Figure S3B) we knew that ICAM-1 was widely expressed on the human ECs and that this expression of this receptor could be increased by pre-treatment with TNFα (Figure 3A). Using *in vitro* static binding assays, we confirmed that the parasite specifically bound to human recombinant ICAM-1 and not the mouse homologue (Figure 3B), confirming that the parasite would only bind to human endothelium and was therefore suitable for our study.

**Figure 3.**
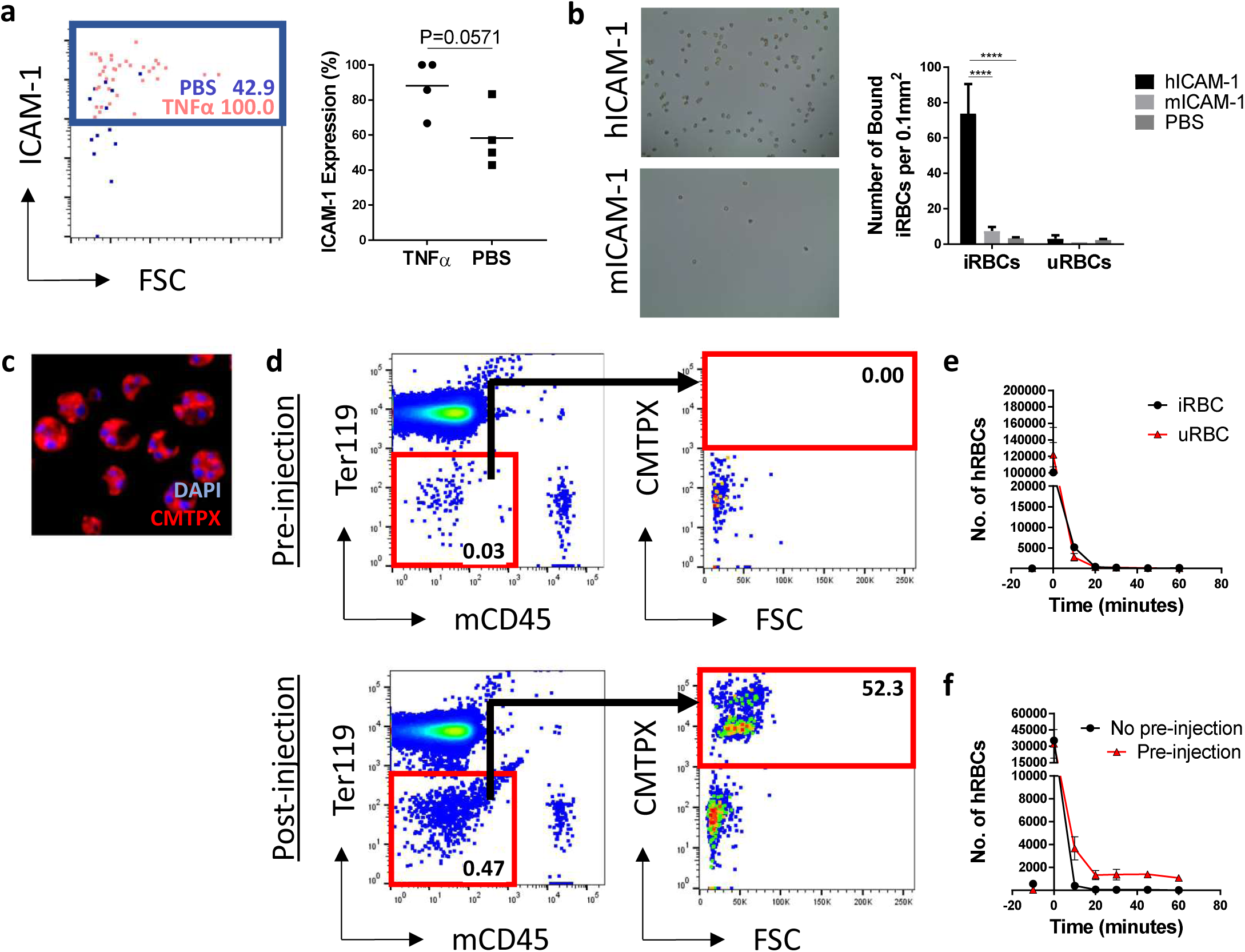
Tracking labelled red blood cells *in vivo*. (a) Human endothelial cell ICAM-1 expression was induced by intravenously injecting engrafted mice with human TNFα (5μg/mouse) 6 hours before they were culled. Control mice received PBS alone. n=4, one independent experiment. Statistical analysis was performed using a Mann-Whitney test, *P<0.05. (b) The *P. falciparum* parasite, ItG, was shown to bind specifically to human recombinant ICAM-1 using *in vitro* binding assays. n=3, one independent experiment. Statistical analysis was performed using a Two-Way ANOVA with Tukey’s multiple comparisons, ****p<0.0001. (c) Infected and uninfected RBCs were labelled with CMTPX. (d-f) RBCs were injected intravenously into mice with human tissue implants and their depletion from the circulation was assessed by flow cytometry. The number of CMPTX labelled uninfected and infected RBCs was assessed without (**e)** and with (**f**) a pre-injection of unlabelled, uninfected RBCs. n=4, one independent experiment.

Parasites were purified in the trophozoite stage of their lifecycle and labelled with the intracellular fluorescent dye, CMTPX (Figure 3C). Following injection, we determined the circulation time of the parasite infected (iRBCs) and uninfected (uRBCs) red blood cells by taking regular blood samples over an hour and quantifying cell number by flow cytometry (Figure 3D). This indicated that the parasites had a short circulation time, being cleared from the blood entirely within 20 minutes of the initial injection (Figure 3E). However, we found that we could extend the circulation time of the labelled cells by giving a pre-injection of 5×10^9^ unlabelled uninfected RBCs to saturate the mouse macrophages that were responsible for removing the cells^27^ (Figure 3F). This allowed approximately 20,000 labelled RBCs to remain in the circulation throughout the imaging period allowing us to perform intravital imaging studies.

### *P. falciparum* cytoadherence can be observed in human blood vessels

After confirming that the *P. falciparum* parasite could bind to human ICAM-1 *in vitro*, we began intravital imaging studies (Figure S5). Prior to imaging labelled RBCs, mice were pre-treated with unlabelled, uninfected RBCs to increase circulation time, TNFα to enhance ICAM-1 expressi on on transplants and finally a fluorescently labelled human anti-CD31 antibody to identify human blood vessels *in vivo*, following the protocols established above. Mice were then injected intravenously with labelled infected RBCs (iRBCs) and a multiphoton microscope was used to image the cells interacting with the labelled human vasculature. Several behaviours associated with cytoadherence were observed including cell tethering and rolling (Figure 4A, Supplementary Video 1), adherence of iRBCs to blood vessel walls (Figure 4B, Supplementary Video 2), blood vessel occlusion (Supplementary Video 2) and collections of iRBC around blood vessel junctions (Figure 4C, Supplementary Video 3). An example of the analysis we could perform on the circulating iRBCs indicated differences in the speed of cells as they sequestered in blood vessels (Figure 4D). Sequestration was confirmed to be specific to human blood vessels through flow cytometry (Figure S6, Figure 4E+4F), which shown a significantly higher number of iRBCs in the implant ear compared to the control ear (Paired T Test, P<0.05).

**Figure 4.**
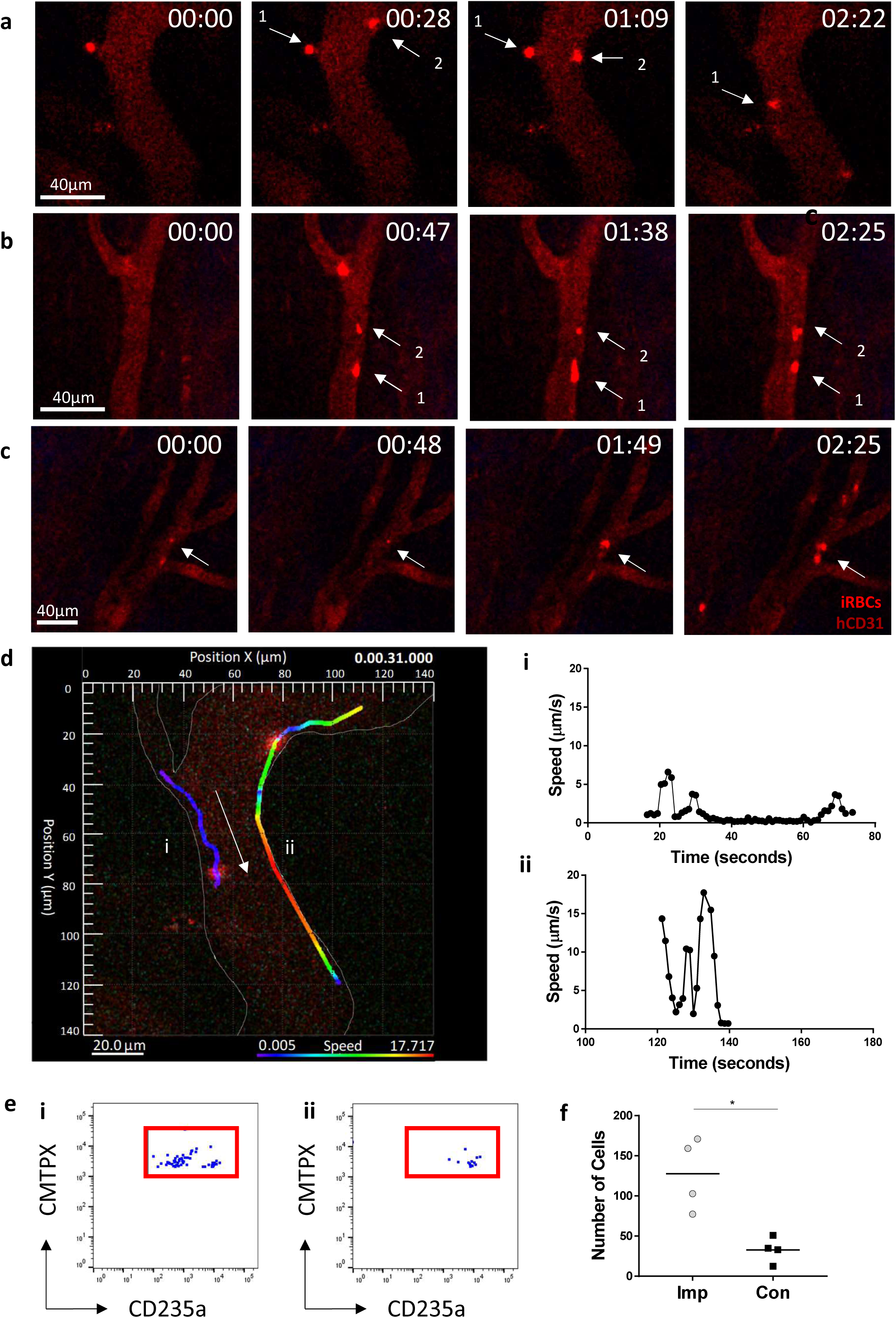
*P. falciparum* cytoadherence can be observed in human blood vessels. Intravital imaging was performed using a multiphoton microscope. Mice were pre-treated with TNFα, unlabelled, uninfected RBCs, and AF647 labelled anti-human CD31 antibody. Labelled iRBCs were then injected into the tail vein of the mice. (a) Tethering and rolling (b) adherence of iRBCs to blood vessel walls (c) and collections of iRBCs around blood vessel junctions were all observed. n=3, three independent experiments. (d) An example of imaging analysis indicated differences in the speed of sequestering cells over time in two different iRBCs (i & ii). Arrow indicates direction of blood flow. Rainbow bar indicates speed of iRBC in blood vessel. Flow cytometry was performed on the (e) (i) implant and (ii) control ears to compare the (f) number of iRBCs in the tissues. n=4, from one independent experiment. Statistical analysis was performed using a paired T test, *p<0.05.

## Discussion

Here, we describe a novel method for culturing and implanting human tissue that promotes the retention and expansion of human vasculature. This system is potentially more biologically relevant than current animal xenograft models, allowing the study and direct observation of human vasculature *in vivo* and the interactions of cells and molecules in vascular processes and diseases. As an example, we demonstrate the application of this approach to the study of *Plasmodium* parasites whose interactions with human vasculature are central to infection associated disease processes.

A key step in maintaining human vasculature in xenografts was to pre-culture tissue in an angiogenic extracellular matrix. This promoted the formation of capillary sprouts and the outgrowth of blood vessels prior to implantation^21^. Previous studies found that this significantly improved the retention of blood vessels of graft origin in murine allografts^22^, however, to the best of our knowledge, this is the first time this has been applied in human xenografts.

Alternative models which employ the use of human endothelial cells lines, such as human umbilical vein endothelial cells (HUVECs), to form microvascular networks have been used extensively to study blood vessel development *in vivo*^26^ but have several drawbacks. These cell lines lack the heterogeneity of ECs and do not account for differences in cell phenotype, which vary greatly depending upon the cells local microenvironment, state of activation and exposure to soluble mediators (Garlanda and Dejana, 1997). Furthermore, the cell lines often lack or have variable levels of many common EC markers including CD34, CD36 and von-Willebrand factor^29,30^ which limits their ability to model the vasculature in different tissues. Our model overcomes these problems by maintaining the grafts endogenous vasculature allowing it to more closely resemble human tissue in its natural state.

Once the grafts were perfused, we noticed few changes in vascularisation between tissues used at different time points post-implantation. This suggested that the vascularisation was physiological and was not persistent and unresolved as is observed with pathological vascularisation. In line with previous studies, the ECs in the grafts were shown to have few phenotypic changes following culture and implantation^31^. However, we did observe a decrease in CD36 expression in implanted tissue, a marker that is particularly enriched in the microvasculature of SAT^32^. The decrease in CD36 expression in the flow cytometry data, alongside the increased vascular diameters suggested a loss of microvasculature in the grafts. We hypothesise that this may be due to changes in the microvascular organisation around the periphery of the graft.

The choice of the ear pinna as an engraftment site had several benefits over other imaging methods, particularly imaging windows^33^. As the ear could be imaged without surgical exposure, we were able to perform intravital imaging of the mice over an extended period of time. This not only reduced the stress that the animal was under but also allowed for the same engrafted animal to be imaged repeatedly, resulting in a significant reduction in the number of animals required for experiments.

Using this method, we chose to examine cytoadherence of the human malarial parasite *P. falciparum* to human endothelium. This is a process that cannot be observed in murine *Plasmodium* infections due to the specific properties of the human-infective parasite^34^. Using our model, we were able to observe the behaviours associated with cytoadherence including rolling, adherence and vessel occlusion^35^. The ItG parasite used in this study was panned on ICAM-1 but was also capable of binding to CD36. As mentioned CD36 expression was downregulated in our ECs following implantation, which may have interfered with the ability of the parasites to sequester in the blood vessels. However, the level of ligand expression on the EC surface required for cytoadherence is unclear. Furthermore, although CD36 is the major adherence receptor in experimental cerebral malaria, it is not clinically significant in the human form of the disease as the adhesion protein on the surface of iRBCs, PfEMP-1, is capable of binding to a large number of receptors and is most associated with endothelial protein C receptor (EPCR) ^36^. This suggests that the impact of reduced CD36 expression would be minimal for studying other parasite strains using this model.

Previous xenograft models^37,38^ have utilised skin in *P. falciparum* research, however, this tissue is not recognised as a particularly significant site of parasite sequestration and cytoadherence^39,40^. By contrast, blood vessels in SAT are a more prominent site of parasite sequestration and previous studies have investigated this tissue as a model to study molecular processes leading to cytoadherence in brain vasculature^41^. For these reasons, we focussed on transplantation of human SAT, however, a range of other tissues were also found to be compatible with the culturing and implantation method. The only tissues we were unable to successfully engraft were those that came from the digestive tract or brain. The ability to transplant a variety of tissues, including SAT, therefore makes our current approach more flexible and better suited to studying the molecular processes of parasite cytoadherence *in vivo*.

It is notoriously difficult to reconstitute human RBCs in mice due to their rapid removal from the circulation by splenic macrophages^27^. In our model, we overcame this issue by giving the mice a pre-injection of uRBCs to saturate the macrophages, allowing us to extend the imaging window for our intravital work. An alternative method would have been to deplete the macrophages in advance using clodronate liposomes^42^; however, due to the severely immunocompromised state of our mice we decided against this method due to welfare concerns.

Although this imaging work is preliminary, the model holds great potential in further assessing the behaviours of parasites, including *P. falciparum*, trypanosomes and schistosomes, as well as bacterial infections such as *Neisseria meningitidis*, which are all known to mediate effects in both the skin and adipose tissue in patients^13,43^. Aside from infection research, the model could also be further adapted to include components of a human immune system, either through the transfer of peripheral blood mononuclear cells (PBMCs) or the transplantation of haematopoietic stem cells, which is a feature often incorporated into humanised models^44^. This would allow us to visualise various aspects of immune cell behaviours, including extravasation, in real time using acute models of inflammation. Other applications in which a model of human vasculature would also be useful include diagnostic imaging in cardiovascular research^45^ and drug screening in cancer studies^12^.

In summary, we have developed a humanised xenograft model in the ear pinna of immunocompromised mice. This model retains the tissues endogenous human vasculature allowing for non-invasive longitudinal imaging of blood flow and cellular recruitment in a system that can be easily adapted to study a variety of different biological processes.

## Author Contributions

GRM designed the research, performed the experiments, constructed the figures, analysed the data and wrote the manuscript. HES provided technical assistance and contributed to experimental design. RO and MDN provided technical assistance and provided input in *Plasmodium* experiments. JCL designed and performed *in vitro* human tissue culture experiments. MM provided assistance in experimental design. PG, AC and JMB designed the research and contributed to writing the paper.

## Acknowledgments

We acknowledge the assistance of both the Institute of Infection, Immunity and Inflammation Flow Core Facility and the Histology Research Service at the University of Glasgow. We also acknowledge the support of NHS Research Scotland (NRS) Greater Glasgow and Clyde Biorepository in acquiring and processing human tissue samples. In addition, we acknowledge Janet Storm for kindly providing a protocol for the static binding assay.

## Data Availability

The datasets generated during and/or analysed during the current study are available from the corresponding author on reasonable request

## References

1. Morton, J. J., Bird, G., Refaeli, Y. & Jimeno, A. Humanized Mouse Xenograft Models⍰: Narrowing the Tumor – Microenvironment Gap. Cancer Res. 76, 6153–6159 (2016).

2. Yong, K., Her, Z. & Chen, Q. Humanized Mice as Unique Tools for Human-Specific Studies. Arch. Immunol. Ther. Exp. (Warsz). 66, 245–266 (2018).

3. Shultz, L. D., Brehm, M. A., Garcia-martinez, J. V. & Greiner, D. L. Humanized mice for immune system investigation⍰: progress, promise and challenges. Nat. Publ. Gr. 12, 786–798 (2012).2370

4. Saunders, M. C., Keller, J. T., Dunsker, S. B. & Mayfield, F. H. Survival of autologous fat grafts in humans and in mice. Connect Tissue Res 8, 85–91 (1981).

5. Geens, M. et al. Spermatogonial survival after grafting human testicular tissue to immunodeficient mice. 21, 390–396 (2006).

6. Holen, I., Speirs, V., Morrissey, B. & Blyth, K. In vivo models in breast cancer research: progress, challenges and future directions. Dis. Model. Mech. 10, 359–371 (2017).

7. Mansour, A. A. et al. An in vivo model of functional and vascularized human brain organoids. Nat. Biotechnol. 36, 432–441 (2018).

8. Colton, C. K. Implantable biohybrid artificial organs. Cell Transplant. 4, 415–436 (1995).

9. Stein, I., Neeman, M., Shweiki, D., Itan, A. & Keshet, E. Stabilization of vascular endothelial growth factor mRNA by hypoxia and hypoglycemia and coregulation with other ischemia-induced genes. Mol. Cell. Biol. 15, 5363–5368 (1995).

10. Calcagni, M. et al. In vivo visualization of the origination of skin graft vasculature in a wild-type/GFP crossover model. Microvasc. Res. 82, 237–245 (2011).

11. Hylander, B. L. et al. Origin of the vasculature supporting growth of primary patient tumor xenografts. J. Transl. Med. 11, 1–14 (2013).

12. Dong, Z. et al. Xenograft tumors vascularized with murine blood vessels may overestimate the effect of anti-tumor drugs: A pilot study. PLoS One 8, 1–8 (2013).

13. De Niz, M. et al. Intravital imaging of host-parasite interactions in skin and adipose tissues. Cell. Microbiol. (2019).

14. Storm, J. & Craig, A. G. Pathogenesis of cerebral malaria — inflammation and cytoadherence MALARIA. Front. Cell. Infect. Microbiol. 4, 1–8 (2014).

15. Ito, M. et al. NOD/SCID/_yc null mouse: an excellent recipient mouse modelfor engraftment of human cells. Blood 100, 3175–3182 (2002).

16. Madkhali, A. M. et al. An Analysis of the Binding Characteristics of a Panel of Recently Selected ICAM-1 Binding Plasmodium falciparum Patient Isolates. PLoS One 9, 4–11 (2014).

17. Trager, W. & Jensen, J. Human Malaria parasites in continuous culture. Science (80-.). 193, 673–676 (1976).

18. Ockenhouse, C. F. et al. Molecular Basis of Sequestration in Severe and Uncomplicated Plasmodium Falciparum Malaria⍰: Differential Adhesion of Infected Erythrocytes to CD36 and ICAM-1. J. Infect 164, 163–169 (1991).

19. Desruisseaux, M. S., Nagajyothi Trujillo, M. E., Tanowitz, H. B. & Scherer, P. E. Adipocyte, adipose tissue, and infectious disease. Infect. Immun. 75, 1066–1078 (2007).

20. Koethe, J. R., Hulgan, T. & Niswender, K. Adipose tissue and immune function: A review of evidence relevant to HIV infection. J. Infect. Dis. 208, 1194–1201 (2013).

21. Rojas-rodriguez, R. et al. Adipose Tissue Angiogenesis Assay. Methods of Adipose Tissue Biology, Part A 537, (Elsevier Inc., 2014).

22. Laschke, M. W. et al. Promoting external inosculation of prevascularised tissue constructs by pre-cultivation in an angiogenic extracellular matrix. Eur. Cells Mater. 20, 356–366 (2010).

23. Gibson, V. B. et al. A novel method to allow noninvasive, longitudinal imaging of the murine immune system in vivo. Blood 119, 1–3 (2012).

24. Swerlick, R. A., Lee, K. H., Wick, T. M. & Lawley, T. J. Human dermal microvascular endothelial but not human umbilical vein endothelial cells express CD36 in vivo and in vitro. J. Immunol. 148, 78–83 (1992).

25. Laschke, M. W. et al. Vascularisation of porous scaffolds is improved by incorporation of adipose tissue-derived microvascular fragments. Eur. Cells Mater. 24, 266–277 (2012).

26. Schechner, J. S. et al. In vivo formation of complex microvessels lined by human endothelial cells in an immunodeficient mouse. PNAS 97, 9191–9196 (2000).

27. Hu, Z., Rooijen, N. V & Yang, Y. Macrophages prevent human red blood cell reconstitution in immunodeficient mice. Blood 118, 5938–5946 (2012).

28. Garlanda, C. & Dejana, E. Heterogeneity of Endothelial Cells. Arterioscler. Thromb. Vasc. Biol. 17, 1193–1202 (1997).

29. McCormick, C. J., Craig, A., Roberts, D., Newbold, C. I. & Berendt, A. R. Intercellular adhesion molecule-1 and CD36 synergize to mediate adherence of Plasmodium falciparum-infected erythrocytes to cultured human microvascular endothelial cells. J. Clin. Invest. 100, 2521–2529 (1997).

30. Müller, A. M. et al. Expression of the endothelial markers PECAM-1, vWF, and CD34 in Vivo and in Vitro. Exp. Mol. Pathol. 72, 221–229 (2002).

31. Motherwell, J. M., Anderson, C. R. & Murfee, W. L. Endothelial Cell Phenotypes are Maintained During Angiogenesis in Cultured Microvascular Networks. Sci. Rep. 8, 1–11 (2018).

32. Silverstein, R. L. & Febbraio, M. CD36, a Scavenger Receptor Involved in Immunity, Metabolism, Angiogenesis, and Behavior. Sci. Signal. 2, 1–8 (2009).

33. Laschke, M. W., Vollmar, B. & Menger, M. D. The dorsal skinfold chamber: Window into the dynamic interaction of biomaterials with their surrounding host tissue. Eur. Cells Mater. 22, 147–167 (2011).

34. Strangward, P. et al. A quantitative brain map of experimental cerebral malaria pathology. PLoS Pathog. 13, 1–37 (2017).

35. Helms, G., Dasanna, A. K., Schwarz, U. S. & Lanzer, M. Modeling cytoadhesion of Plasmodium falciparum-infected erythrocytes and leukocytes—common principles and distinctive features. FEBS Lett. 1955–1971 (2016). doi:10.1002/1873-3468.12142

36. Mkumbaye, S. I. et al. The Severity of Plasmodium falciparum Infection Is Associated with Transcript Levels of var Genes Encoding Endothelial Protein C Receptor-Binding P. falciparum Erythrocyte Membrane Protein 1. Cell. Microbiol. 85, 1–14 (2017).

37. Ho, M., Hickey, M. J., Murray, A. G., Andonegui, G. & Kubes, P. Visualization of Plasmodium falciparum–Endothelium Interactions in Human Microvasculature: Mimicry of Leukocyte Recruitment. J. Exp. Med. 192, 1205–1212 (2000).

38. Yipp, B. G. et al. Differential roles of CD36, ICAM-1, and p-selectin in Plasmodium falciparum cytoadherence in vivo. Microcirculation 14, 593–602 (2007).

39. Jervis, H. R., Sprinz, H., Johnson, A. J. & Wellde, A. T. Experimental Infection in With. Am. J. Trop. Med. Hygeine 21, 382–386 (1972).

40. MacPherson, G. G., Warrell, M. J., White, N. J., Looareesuwan, S. & Warrell, D. A. Human cerebral malaria. A quantitative ultrastructural analysis of parasitized erythrocyte sequestration. Am. J. Pathol. 119, 385–401 (1985).

41. Moxon, C. A. et al. Loss of endothelial protein C receptors links coagulation and inflammation to parasite sequestration in cerebral malaria in African children. Blood 122, 842–852 (2013).

42. Amaladoss, A. et al. De Novo Generated Human Red Blood Cells in Humanized Mice Support Plasmodium falciparum Infection. PLoS One 10, e0129825 (2015).

43. Bruneval, P., Duménil, G., Michea Veloso, P., Martin, T. & Melican, K. Adhesion of Neisseria meningitidis to Dermal Vessels Leads to Local Vascular Damage and Purpura in a Humanized Mouse Model. PLoS Pathog. 9, e1003139 (2013).

44. Minkah, N. K., Schafer, C. & Kappe, S. H. I. Humanized Mouse Models for the Study of Human Malaria Parasite Biology, Pathogenesis, and immunity. Front. Immunol. 9, (2018).

45. Noonan, J. et al. In vivo multiplex molecular imaging of vascular inflammation using surface-enhanced Raman spectroscopy. Theranostics 8, 6195–6209 (2018).

