## Supplementary Figure Legends for "Developing a xenograft model of human vasculature in the mouse ear pinna"

**Supplementary Figure 1 – Alternative tissues can be implanted into the ear pinna.**

Human (i) dermis, (ii) kidney cortex, (iii) skeletal muscle, (iv) brain, (v) duodenum and (vi) colon were implanted into the ear pinna of immunocompromised mice. The mice were culled three weeks post implantation, and their ears were sectioned and processed for histology.

**Supplementary Figure 2 – Gating strategy for identifying endothelial cells by flow cytometry**

Cells were gated on FSC/SSC, single cells, viable cells, mCD45-/hCD45-, mCD31+ or hCD31+

**Supplementary Figure 3 - Changes in expression of endothelial cells markers**

The expression of the endothelial cell markers (a) CD34, (b) ICAM-1 and (c) CD36 were assessed by flow cytometry. n=5 from five independent experiments. Each data point from implanted samples was pooled from 3-4 mice implanted with human tissue from the same source. Statistical analysis was performed using a Two-Way ANOVA, *p<0.05, ***p<0.001, ****p<0.0001.

**Supplementary Figure 4 – 3D Image of Engrafted Adipose Tissue**

A 3D image was taken of engrafted adipose tissue following an intravenous injection of an anti-human CD31 antibody.

**Supplementary Figure 5 – Intravital Imaging of Engrafted Mice**

Mice were imaged intravitally using a (i) multiphoton microscope. (ii) Mice were first placed on a metal stage and anaesthetised with a face mask. (iii) The ear was then affixed to the stage using veterinary glue and a (iv) silicone grease ring was made around the ear and filled with PBS. (v) The objective was then lowered, allowing the imaging studies to be performed.

**Supplementary Figure 6 – Gating strategy for identifying labelled human RBCs in mouse tissue**

Cells were gated on FSC/SSC, single cells, V450-/V500- (to exclude autofluorescence), Ter119-/mCD45- (to exclude murine RBCs/leukocytes), CMTPX+, CMTPX+/CD235a (human RBCs)
