## Supplementary Figures for "Developing a xenograft model of human vasculature in the mouse ear pinna"

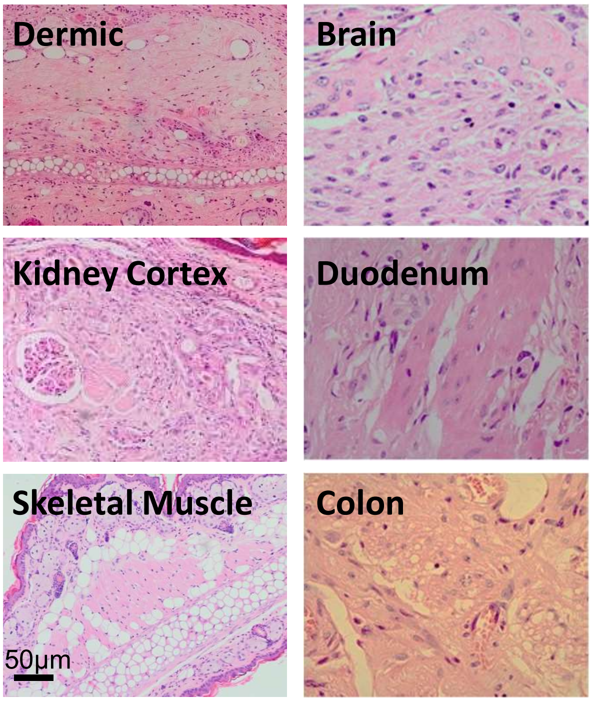


**Supplementary Figure 1 – Alternative tissues can be implanted into the ear pinna.**

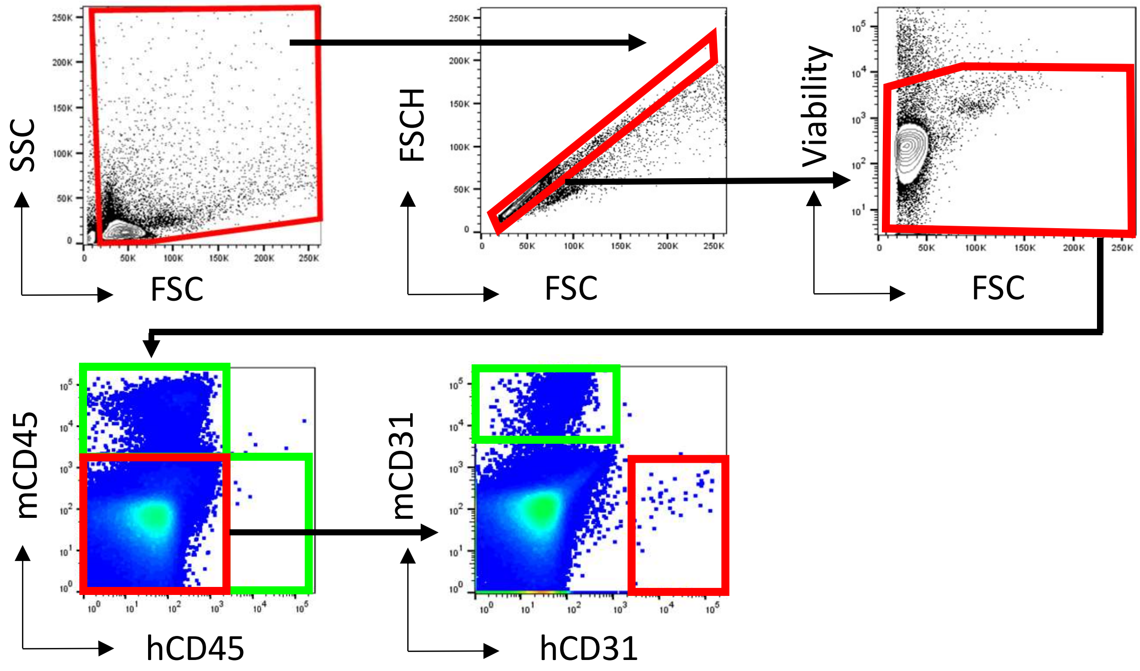


**Supplementary Figure 2 – Gating strategy for identifying endothelial cells by flow cytometry**

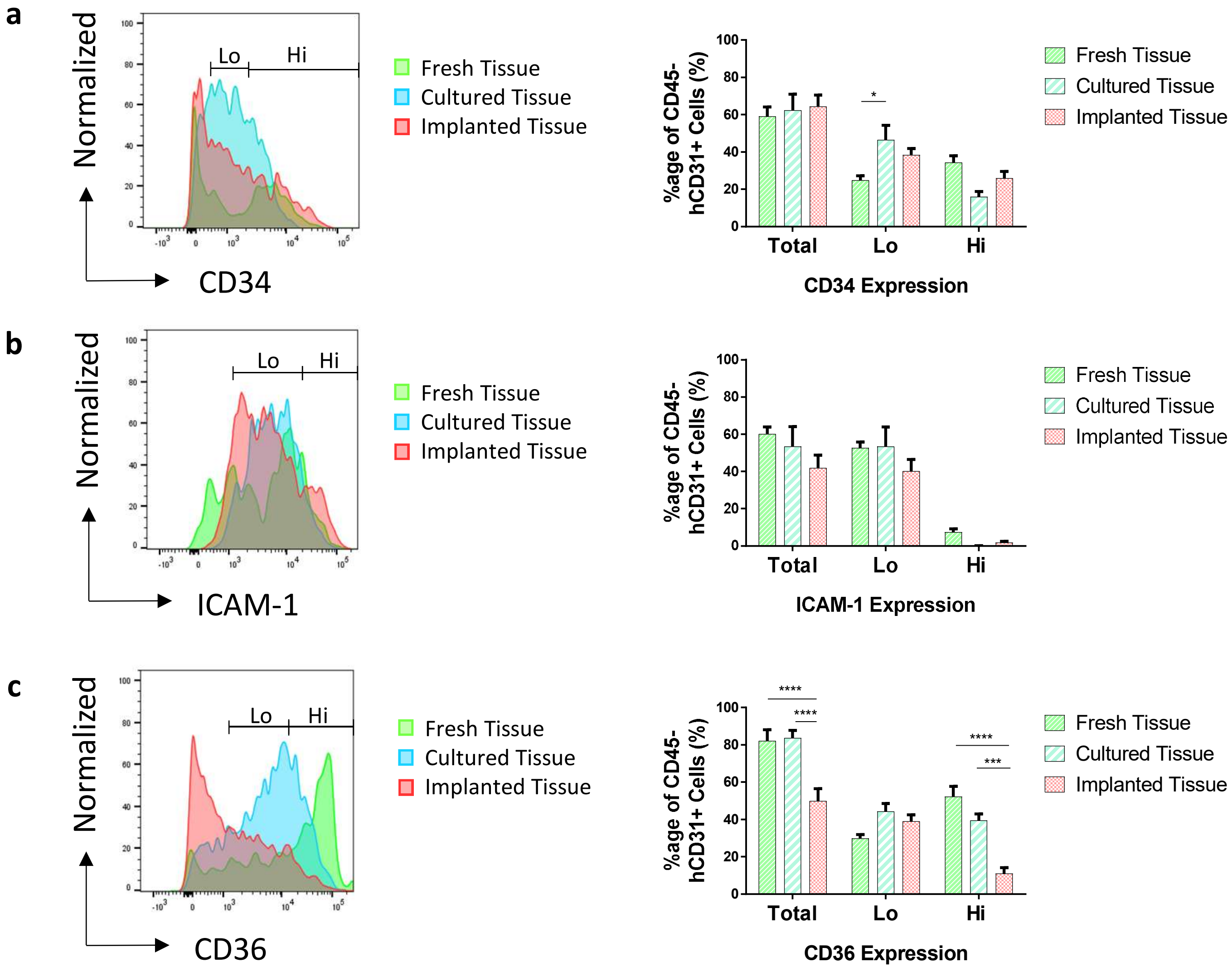


**Supplementary Figure 3 - Changes in expression of endothelial cells markers**

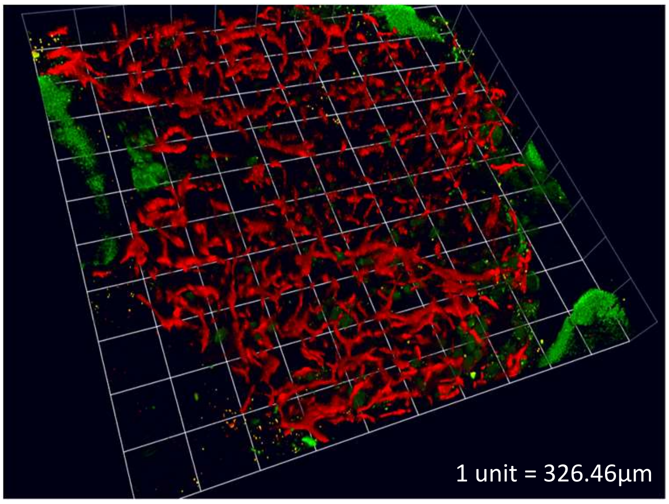


**Supplementary Figure 4 – 3D Image of engrafted adipose tissue**

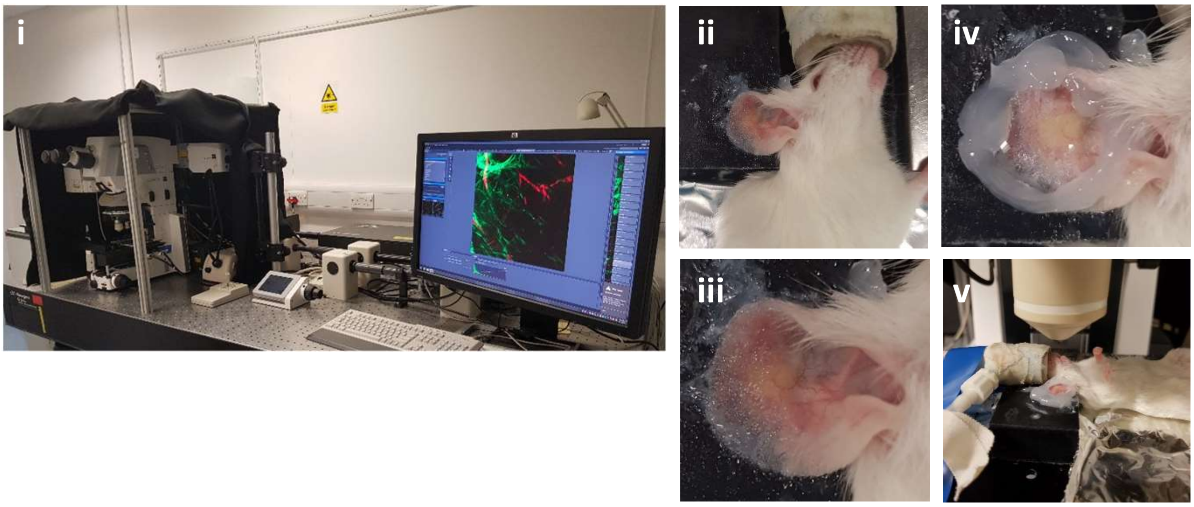


**Supplementary Figure 5 – Intravital imaging of engrafted mice**

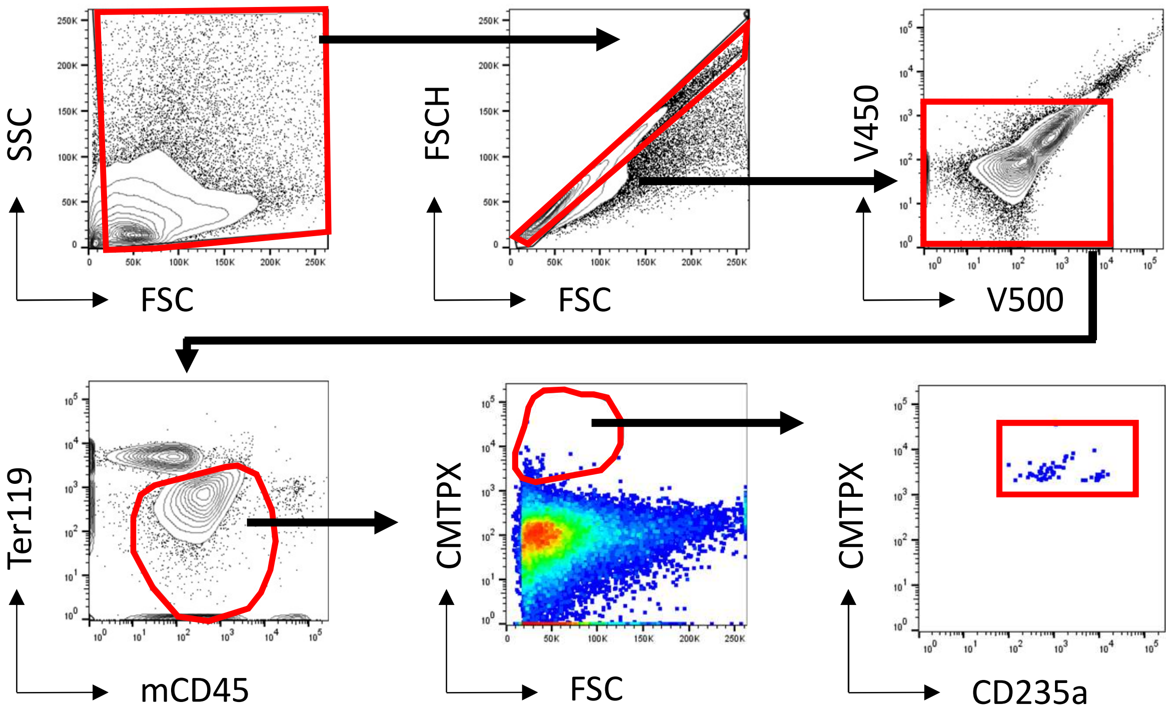


**Supplementary Figure 6 – Gating strategy for identifying labelled human RBCs in mouse tissue**
